# Viral fragments in the urine proteome: New clues to the cause of fever

**DOI:** 10.1101/2024.09.19.613984

**Authors:** Minhui Yang, Chenyang Zhao, Youhe Gao

## Abstract

**[Background]:** Fever of unknown origin refers to a medical condition where the cause of the fever is not yet clear. This condition is common in various potential diseases and usually requires detailed examination and testing to determine the specific cause. No one has ever looked for the cause of fever in urine proteomics, and this study provides clues and diagnostic evidence for patients with fever of unknown origin through urine proteomics analysis.

**[Methods]:** We attempted virus-wide database searching for the first time in urine samples, using liquid chromatography-tandem mass spectrometry (LC-MS/MS) to analyze urine proteins from febrile patients. Virus protein fragments were identified and retrieved.

**[Results]:** In urine samples, we detected specific peptide segments from various viruses including monkeypox virus, saliva virus A, human herpesvirus 8 type P, Middle East respiratory syndrome-related coronavirus, rotavirus A, foot-and-mouth disease virus (strain NZ2), human herpesvirus 2 (strain HG52), human adenovirus E serotype 4, influenza A virus, human coronavirus NL63, influenza B virus (strain W3), Nipah virus, and hepatitis C virus genotype 2k (isolate VAT96), among others. Several viruses showed significantly higher detection levels in febrile patients compared to controls, with saliva virus A showing an increase by over 4200-fold, and multiple virus proteins increased by more than 20-fold. It is noteworthy that the overall protein retrieval error rate was less than 1%, and the individual protein retrieval error rate for each sample was less than 5%, ensuring highly reliable protein retrieval results with a probability of error across all proteins of only 1.2×e-17.

**[Conclusion]:** Reliable virus protein fragments can be retrieved from urine proteomics, providing clues for febrile patient investigation and potentially applicable to the exploration of any unknown diseases. By adopting our method, there is no need to know in advance what specific viruses are contained in the sample, as long as the information of these viruses has been stored in the virus database, comprehensive and extensive virus retrieval can be achieved. This method significantly expands the coverage of virus detection and improves the flexibility and efficiency of detection.

## Introduction

Fever, as a non-specific response of the body to potential health threats, may hide a variety of causes, such as bacterial infection, viral infection, fungal infection, parasitic infection, as well as inflammation, drug side effects and tumors ^[1]^. In view of the various causes of fever, it is particularly important for doctors to accurately diagnose the exact causes behind it in the process of diagnosis and treatment. The accuracy of this diagnosis is not only related to the health recovery of patients, but also the key to the effective implementation of follow-up treatment.

Rapid identification of the exact cause of fever is not only helpful to ensure the accurate diagnosis of the disease, but also to start the treatment process in time, effectively curb the spread of the disease, improve treatment efficiency, and reduce medical costs. Through timely and accurate diagnosis, doctors can formulate targeted treatment plans to avoid unnecessary treatment and drug use, so as to protect patients’ health and improve treatment effect ^[2]^. Urine proteomics has shown great potential in the exploration and analysis of biomarkers. Compared with other biological samples, urine has unique advantages: it is less strictly regulated by the physiological steady-state mechanism, so it can more sensitively capture the subtle biochemical fluctuations in the body. In addition, urine can be obtained without professional collection means, and the collection process is not only non-invasive, but also convenient and fast, which makes it an ideal sample source for biomarker research ^[3, 4]^. In recent years, studies have proved that urine detection has potential in virus monitoring and diagnosis. For example, during the covid-19 epidemic, scientists successfully detected the genetic material of neocoronavirus in the urine of patients, which provided the possibility of non-invasive and simple diagnostic methods. Similarly, the genetic material of West Nile virus was also found in urine ^[5]^. In addition, some scientists also detected sars-cov-2 nucleocapsid protein derived peptide in urine samples ^[6]^. These technologies have shown broad application prospects. Therefore, as a non-invasive and repeatable sampling method, urine detection is of great significance for the monitoring and early diagnosis of viral diseases, and is expected to play an increasingly important role in clinical practice.

In this study, we searched the full viral Library of urine proteins of fever patients and normal people, and analyzed them in fever group and normal group. At the same time, because the causes of fever in each patient may be different, we also compared each patient with the normal group, in order to provide clues and basis for the clinical diagnosis of fever through the differential proteins of virus.

## 1 Material and method

### 1.1 collection of patient samples

The experimental materials were collected by zhaochenyang in the laboratory. The details are as follows: 11 urine samples from patients with fever at admission were collected from Beijing China Japan Friendship Hospital, all of which were morning urine. The specific information is shown in Table 1. These samples were obtained in the laboratory department, without any patient identity information, and did not affect any treatment or recommend any clinical and auxiliary examinations. The purpose of the study is to obtain clues of fever, and the knowledge obtained is only used for research purposes. The patient’s temperature record is shown in Table 1. All participants have signed the informed consent, and the study has been ethically approved by the China Japan Friendship Hospital. (No.: 2019-42-k30).

**Table1.** Records of clinical characteristics of patients.

| Patient number | Sex | Age | Body temperature /°C |
| --- | --- | --- | --- |
| F1 | female | 60 | 38.8 |
| F2 | female | 81 | 38.4 |
| F3 | male | 60 | 37.3 |
| F4 | male | 73 | 37.8 |
| F5 | male | 72 | 36.0 |
| F6 | male | 64 | 36.6 |
| F7 | male | 32 | 36.5 |
| F8 | male | 71 | 36.8 |
| F9 | female | 53 | 36.6 |
| F10 | male | 88 | 36.8 |
| F11 | female | 32 | 36.9 |

### 1.2 Processing of urine samples

The collected urine samples are stored in the -80°C refrigerator for future use. First, after defrosting the 4ml urine sample, the cell debris was removed by centrifugal force at 12000×g at 4°C for 20 minutes. Then, the supernatant was mixed with 20 mM DTT and heated in a metal bath at 99.2°C for 10 minutes. After cooling to room temperature, 50 mM IAA was added for light shielding reaction for 40 minutes. After the reaction, the sample was fully mixed with four times the volume of pre-cooled anhydrous ethanol and placed in a -20°C refrigerator to precipitate the protein for 24 hours. After precipitation, centrifuge at 4°C, 10000×g for 30 min, remove the supernatant, blow dry the protein precipitation, and add appropriate lysate (8 mol/L urea, 2 mol/L thiourea, 50 mmol/L Tris and 25 mmol/L DTT) to redissolve. The redissolved samples were centrifuged, the supernatant was retained, and the protein concentration was determined by Bradford method.

Urinary protease cutting: The auxiliary enzyme cutting on the urinary protein membrane was performed by FASP method. 100ug of urine protein was added to the filter membrane of 10kD ultrafiltration tube, and washed twice with UA solution of 8 mol/L urea and 0.1 mol/L Tris-HCl (pH8.5) and 25 mmol/L NH4HCO3 solution, respectively. Subsequently, trypsin was added at a ratio of 1:50 trypsin: protein and incubated at 37°C overnight. After overnight, the filtrate after enzymatic hydrolysis is collected by centrifugation, which is the polypeptide mixture. Finally, Oasis HLB solid phase extraction column was demineralized, vacuum dried and stored at -80°C.

### 1.3 LC-MS/MS tandem mass spectrometry analysis

The peptide was redissolved with 0.1% formic acid water, and the peptide concentration was diluted to 0.5 μ g/μ L. 1 μ g peptide sample was separated by thermo esay-nlc1200 liquid phase system. The parameters were set as follows: elution time 90 minutes, elution gradient (phase A: 0.1% formic acid; phase B: 80% acetonitrile). The isolated peptides were detected by Orbitrap fusion Lumos tribird mass spectrometer, and data independent acquisition mode was adopted.

### 1.4 Data Analysis

Each polypeptide sample was taken for mass spectrometry in DIA mode, followed by data processing and analysis using Spectronaut X software. We downloaded the latest whole-virus protein data from UniProt to build the database, and then conducted a virus database search on the DIA raw data files of each sample. In this process, the parameters are set as follows: the FDR of the protein is 1%, and the FDR of the peptide is 1%.

## 2. Results and discussion

After the urine samples were processed, 19 protein samples were analyzed by LC-MS/MS tandem mass spectrometry. A total of 26 viral proteins were identified, including 13 specific peptides. It should be noted that the retrieval error rate of total protein is less than 1%, and the retrieval error rate of single protein in each sample is less than 5%. Therefore, our protein retrieval results are very reliable, and the probability of all protein retrieval errors is only 1.2 × E-17. Table 2 shows the viral proteins with specific peptides detected.

**Table2.**
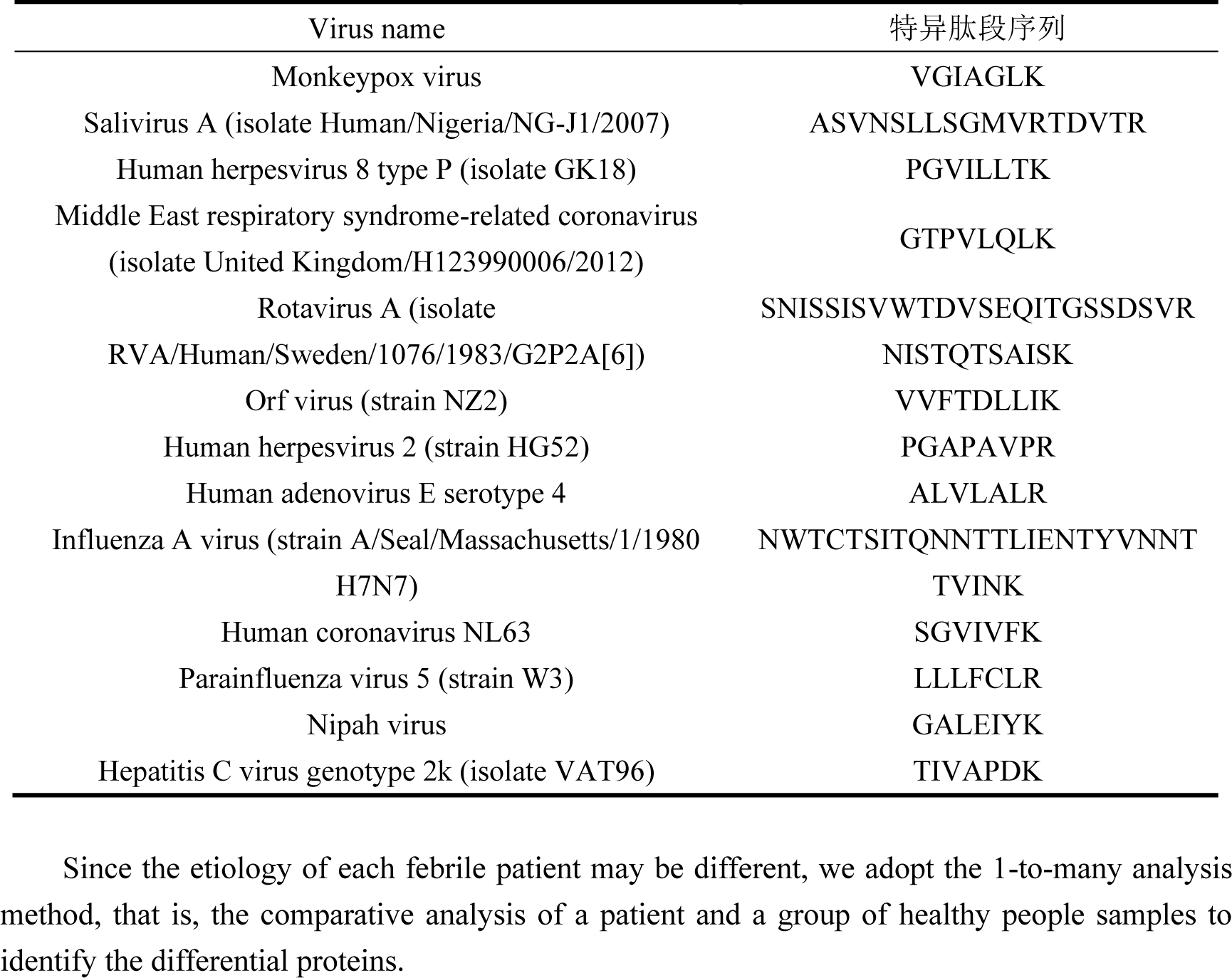
Names and sequences of specific viruses.

### 2.1 Analysis of urinary proteome in single patient

#### (1) Analysis of urinary proteome in F1 patients

Comparing the samples of F1 patients with 8 healthy samples, the detection amount of salivary virus A was 4289 times higher than that of the control group, and the detection amount of rotavirus a was 16 times higher than that of the control group, suggesting that the content of salivary virus and rotavirus a in the patient was higher than that of the normal people, which may be the cause of fever.

**Table3.**
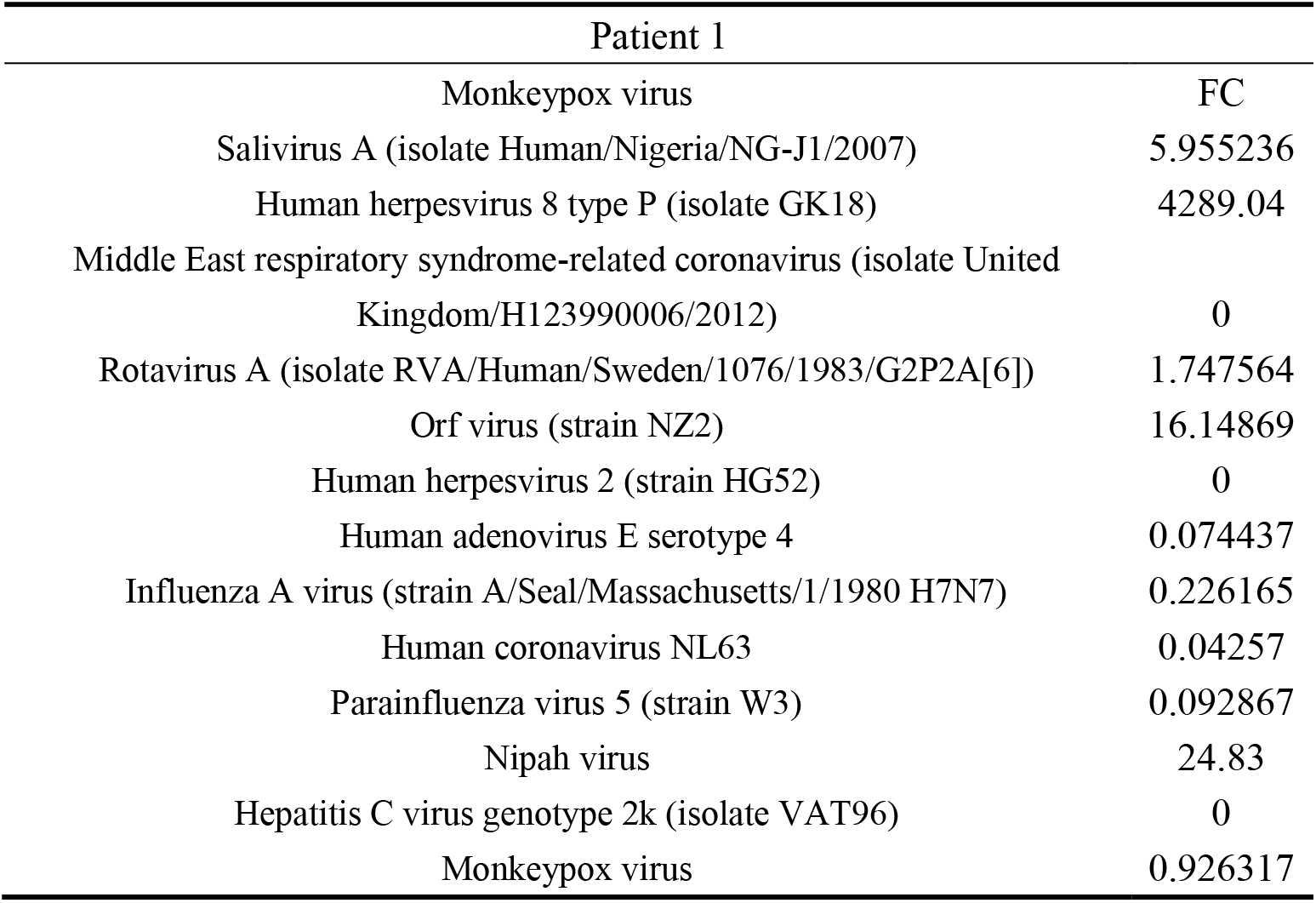
Urine protein analysis of F1 patients.

| Patient 1 |  |
| --- | --- |
| Monkeypox virus | FC |
| Salivirus A (isolate Human/Nigeria/NG-J1/2007) | 5.955236 |
| Human herpesvirus 8 type P (isolate GK18) | 4289.04 |
| Middle East respiratory syndrome-related coronavirus (isolate United Kingdom/H123990006/2012) | 0 |
| Rotavirus A (isolate RVA/Human/Sweden/1076/1983/G2P2A[6]) | 1.747564 |
| Orf virus (strain NZ2) | 16.14869 |
| Human herpesvirus 2 (strain HG52) | 0 |
| Human adenovirus E serotype 4 | 0.074437 |
| Influenza A virus (strain A/Seal/Massachusetts/1/1980 H7N7) | 0.226165 |
| Human coronavirus NL63 | 0.04257 |
| Parainfluenza virus 5 (strain W3) | 0.092867 |
| Nipah virus | 24.83 |
| Hepatitis C virus genotype 2k (isolate VAT96) | 0 |
| Monkeypox virus | 0.926317 |

#### (2) Analysis of urinary proteome in F2 patients

Comparing the samples of F2 patients with 8 healthy samples, the detection amount of salivary virus A was 2962 times higher than that of the control group, and the detection amount of rotavirus a was 20 times higher than that of the control group, as shown in Fig. 3 and Fig. 4, reminding that the content of salivary virus and rotavirus a in the patient was higher than that of the normal person, which may be the cause of fever.

**Table4.** Urine protein analysis of F2 patients.

| Patient 2 |  |
| --- | --- |
| Monkeypox virus | FC |
| Salivirus A (isolate Human/Nigeria/NG-J1/2007) | 8.270417 |
| Human herpesvirus 8 type P (isolate GK18) | 2962.705 |
| Middle East respiratory syndrome-related coronavirus (isolate United Kingdom/H123990006/2012) | 0 |
| Rotavirus A (isolate RVA/Human/Sweden/1076/1983/G2P2A[6]) | 1.451347 |
| Orf virus (strain NZ2) | 20.10132 |
| Human herpesvirus 2 (strain HG52) | 0 |
| Human adenovirus E serotype 4 | 0.065635 |
| Influenza A virus (strain A/Seal/Massachusetts/1/1980 H7N7) | 0.326977 |
| Human coronavirus NL63 | 1.174532 |
| Parainfluenza virus 5 (strain W3) | 0.063747 |
| Nipah virus | 15.67503 |
| Hepatitis C virus genotype 2k (isolate VAT96) | 0 |
| Monkeypox virus | 0.568211 |

#### (3) Analysis of urinary proteome in F3 patients

Comparing the samples of F3 patients with 8 healthy samples, the detection amount of salivary virus A was 18 times higher than that of the control group, and the detection amount of monkeypox virus was 27 times higher than that of the control group, suggesting that the content of salivary virus A and monkeypox virus A in the patient was higher than that of the normal people, which may be the cause of fever.

**Table5.** Urine protein analysis of F3 patients.

| Patient 3 |  |
| --- | --- |
| Monkeypox virus | FC |
| Salivirus A (isolate Human/Nigeria/NG-J1/2007) | 27.39205 |
| Human herpesvirus 8 type P (isolate GK18) | 18.02841 |
| Middle East respiratory syndrome-related coronavirus (isolate United Kingdom/H123990006/2012) | 0 |
| Rotavirus A (isolate RVA/Human/Sweden/1076/1983/G2P2A[6]) | 0.803686 |
| Orf virus (strain NZ2) | 0.382166 |
| Human herpesvirus 2 (strain HG52) | 3.2059 |
| Human adenovirus E serotype 4 | 0.301416 |
| Influenza A virus (strain A/Seal/Massachusetts/1/1980 H7N7) | 0.908352 |
| Human coronavirus NL63 | 3.578544 |
| Parainfluenza virus 5 (strain W3) | 0.173418 |
| Nipah virus | 0.483941 |
| Hepatitis C virus genotype 2k (isolate VAT96) | 0.690899 |
| Monkeypox virus | 0.737359 |

#### (4) Analysis of urinary proteome in F4 patients

Comparing the samples of F4 patients with 8 healthy samples, the detection amount of monkeypox virus was 7 times higher than that of the control group, suggesting that the content of monkeypox virus in the patient was higher than that of the normal people, which may be the cause of fever.

**Table6.** Urine protein analysis of F4 patients.

| Patient 4 |  |
| --- | --- |
| Monkeypox virus | FC |
| Salivirus A (isolate Human/Nigeria/NG-J1/2007) | 7.654227 |
| Human herpesvirus 8 type P (isolate GK18) | 3.254331 |
| Middle East respiratory syndrome-related coronavirus (isolate United Kingdom/H123990006/2012) | 1.020293 |
| Rotavirus A (isolate RVA/Human/Sweden/1076/1983/G2P2A[6]) | 1.610903 |
| Orf virus (strain NZ2) | 2.191586 |
| Human herpesvirus 2 (strain HG52) | 3.200291 |
| Human adenovirus E serotype 4 | 0.386626 |
| Influenza A virus (strain A/Seal/Massachusetts/1/1980 H7N7) | 0.416387 |
| Human coronavirus NL63 | 1.235479 |
| Parainfluenza virus 5 (strain W3) | 0.110488 |
| Nipah virus | 0 |
| Hepatitis C virus genotype 2k (isolate VAT96) | 1.105003 |
| Monkeypox virus | 0.588676 |

#### (5) Analysis of urinary proteome in F5 patients

Comparing the samples of F5 patients with 8 healthy samples, the detection amount of monkeypox virus was 6.9 times higher than that of the control group, suggesting that the content of monkeypox virus in the patient was higher than that of the normal people, which may be the cause of fever.

**Table7.**
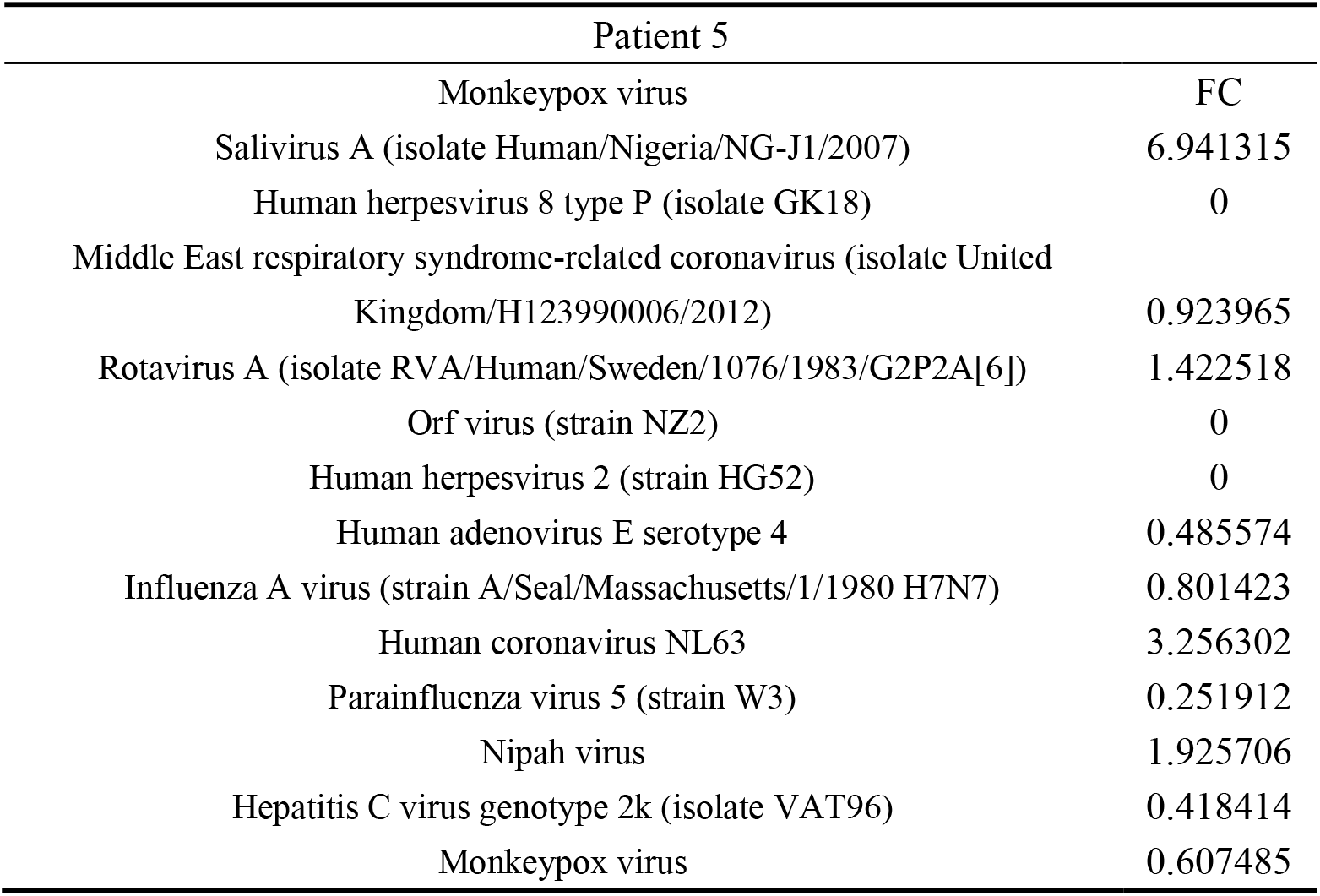
Urine protein analysis of F5 patients.

| Patient 5 |  |
| --- | --- |
| Monkeypox virus | FC |
| Salivirus A (isolate Human/Nigeria/NG-J1/2007) | 6.941315 |
| Human herpesvirus 8 type P (isolate GK18) | 0 |
| Middle East respiratory syndrome-related coronavirus (isolate United Kingdom/H123990006/2012) | 0.923965 |
| Rotavirus A (isolate RVA/Human/Sweden/1076/1983/G2P2A[6]) | 1.422518 |
| Orf virus (strain NZ2) | 0 |
| Human herpesvirus 2 (strain HG52) | 0 |
| Human adenovirus E serotype 4 | 0.485574 |
| Influenza A virus (strain A/Seal/Massachusetts/1/1980 H7N7) | 0.801423 |
| Human coronavirus NL63 | 3.256302 |
| Parainfluenza virus 5 (strain W3) | 0.251912 |
| Nipah virus | 1.925706 |
| Hepatitis C virus genotype 2k (isolate VAT96) | 0.418414 |
| Monkeypox virus | 0.607485 |

#### (6) Analysis of urinary proteome in F6 patients

Comparing the samples of F6 patients with 8 healthy samples, the detection amount of monkeypox virus was 21 times higher than that of the control group, the detection amount of salivary virus A was 77 times higher than that of the control group, and the detection amount of parainfluenza virus 5 was 16 times higher than that of the control group, suggesting that the content of monkeypox virus in the patient was higher than that of the normal person, which may be the cause of fever.

**Table8.** Urine protein analysis of F6 patients.

| Patient 6 |  |
| --- | --- |
| Monkeypox virus | FC |
| Salivirus A (isolate Human/Nigeria/NG-J1/2007) | 21.88945 |
| Human herpesvirus 8 type P (isolate GK18) | 77.49099 |
| Middle East respiratory syndrome-related coronavirus (isolate United Kingdom/H123990006/2012) | 0 |
| Rotavirus A (isolate RVA/Human/Sweden/1076/1983/G2P2A[6]) | 5.259096 |
| Orf virus (strain NZ2) | 0 |
| Human herpesvirus 2 (strain HG52) | 0 |
| Human adenovirus E serotype 4 | 0.446669 |
| Influenza A virus (strain A/Seal/Massachusetts/1/1980 H7N7) | 0.789937 |
| Human coronavirus NL63 | 2.622181 |
| Parainfluenza virus 5 (strain W3) | 0.258942 |
| Nipah virus | 16.54252 |
| Hepatitis C virus genotype 2k (isolate VAT96) | 0.299907 |
| Monkeypox virus | 0.656443 |

#### (7) Analysis of urinary proteome in F7 patients

Comparing the F7 patients’ samples with 8 healthy samples, no significant increase in viral protein was found.

**Table9.** Urine protein analysis of F7 patients.

| Patient 7 |  |
| --- | --- |
| Monkeypox virus | FC |
| Salivirus A (isolate Human/Nigeria/NG-J1/2007) | 0.665584 |
| Human herpesvirus 8 type P (isolate GK18) | 0.105637 |
| Middle East respiratory syndrome-related coronavirus (isolate United Kingdom/H123990006/2012) | 0.873446 |
| Rotavirus A (isolate RVA/Human/Sweden/1076/1983/G2P2A[6]) | 1.323761 |
| Orf virus (strain NZ2) | 0 |
| Human herpesvirus 2 (strain HG52) | 0 |
| Human adenovirus E serotype 4 | 0.388129 |
| Influenza A virus (strain A/Seal/Massachusetts/1/1980 H7N7) | 0.712458 |
| Human coronavirus NL63 | 8.049586 |
| Parainfluenza virus 5 (strain W3) | 0 |
| Nipah virus | 0 |
| Hepatitis C virus genotype 2k (isolate VAT96) | 2.066986 |
| Monkeypox virus | 1.372666 |

#### (8) Analysis of urinary proteome in F8 patients

Comparing F8 patients’ samples with 8 healthy samples, the detection amount of foot-and-mouth disease virus was 10 times higher than that of the control group, suggesting that the content of foot-and-mouth disease virus in the patient was higher than that of the normal person, which may be the cause of fever.

**Table10.**
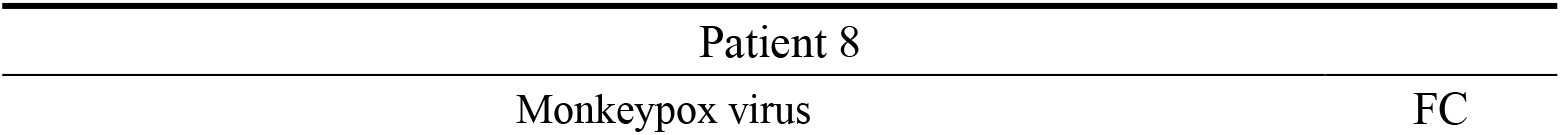
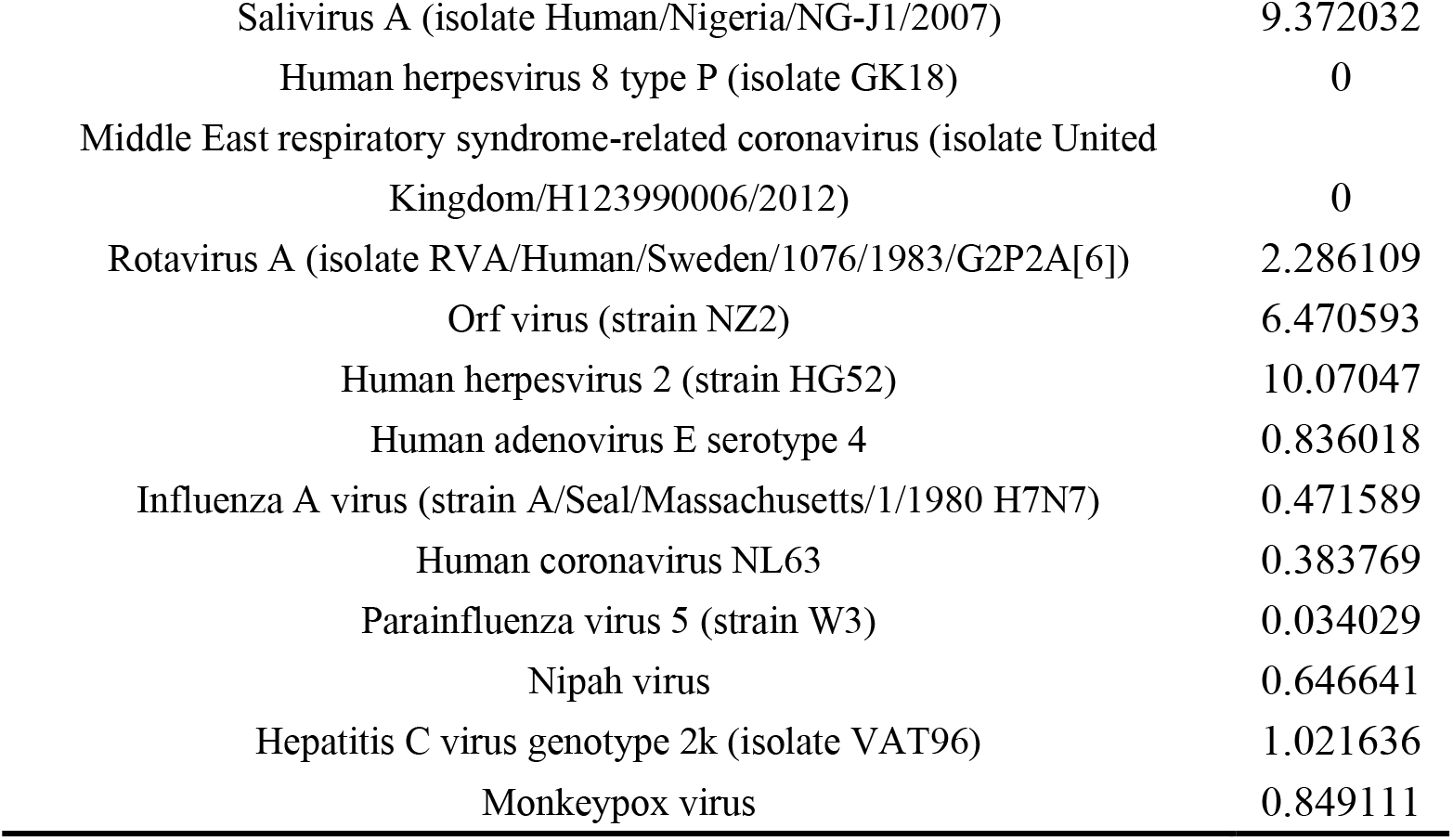
Urine protein analysis of F8 patients.

| Patient 8 |  |
| --- | --- |
| Monkeypox virus | FC |
| Salivirus A (isolate Human/Nigeria/NG-J1/2007) | 9.372032 |
| Human herpesvirus 8 type P (isolate GK18) | 0 |
| Middle East respiratory syndrome-related coronavirus (isolate United Kingdom/H123990006/2012) | 0 |
| Rotavirus A (isolate RVA/Human/Sweden/1076/1983/G2P2A[6]) | 2.286109 |
| Orf virus (strain NZ2) | 6.470593 |
| Human herpesvirus 2 (strain HG52) | 10.07047 |
| Human adenovirus E serotype 4 | 0.836018 |
| Influenza A virus (strain A/Seal/Massachusetts/1/1980 H7N7) | 0.471589 |
| Human coronavirus NL63 | 0.383769 |
| Parainfluenza virus 5 (strain W3) | 0.034029 |
| Nipah virus | 0.646641 |
| Hepatitis C virus genotype 2k (isolate VAT96) | 1.021636 |
| Monkeypox virus | 0.849111 |

#### (9) Analysis of urinary proteome in F9 patients

Comparing the F9 patients’ samples with 8 healthy samples, no significant increase in viral protein was found.

**Table11.** Urine protein analysis of F9 patients.

| Patient 9 |  |
| --- | --- |
| Monkeypox virus | FC |
| Salivirus A (isolate Human/Nigeria/NG-J1/2007) | 4.495728 |
| Human herpesvirus 8 type P (isolate GK18) | 0 |
| Middle East respiratory syndrome-related coronavirus (isolate United Kingdom/H123990006/2012) | 0.487841 |
| Rotavirus A (isolate RVA/Human/Sweden/1076/1983/G2P2A[6]) | 2.226839 |
| Orf virus (strain NZ2) | 0 |
| Human herpesvirus 2 (strain HG52) | 0 |
| Human adenovirus E serotype 4 | 0.331033 |
| Influenza A virus (strain A/Seal/Massachusetts/1/1980 H7N7) | 1.590838 |
| Human coronavirus NL63 | 0.568287 |
| Parainfluenza virus 5 (strain W3) | 0.195666 |
| Nipah virus | 1.006321 |
| Hepatitis C virus genotype 2k (isolate VAT96) | 0.25944 |
| Monkeypox virus | 0.498966 |

By combining urine proteomics and one to many analysis, we found that there were obvious characteristics in the patients with fever to be examined: the amount of virus A and rotavirus a detected in the saliva of patients 1 and 2 was significantly higher than that of the control group; The amount of virus A and monkeypox virus in the saliva of patient 3 also increased significantly; Patients 4 and 5 showed significant increase of monkeypox virus; However, the saliva of patient No. 6 not only detected virus A and monkeypox virus, but also detected parainfluenza virus 5, and their detection levels were significantly higher than those of the control group; Finally, foot and mouth disease virus was detected in the saliva of patient No. 8, and the magnitude was significantly higher than that of the control group.

## 3. Conclusion and Discussion

In this study, liquid chromatography tandem mass spectrometry (LC-MS/MS) technology was used to analyze the urine protein of patients with fever, and a comprehensive virus library search was carried out. The results showed that a variety of viral protein fragments were found in urine proteins, and the content of these protein fragments was significantly different from that of healthy people, such as salivary virus A, foot-and-mouth disease virus, monkeypox virus, parainfluenza virus 5 and other related proteins. This study confirmed the presence of viral protein in urine protein, and provided new clues and ideas for the discovery of fever cases and other viral infectious diseases.

Some viruses can release abundant protein peptides in urine. In order to effectively identify and track these emerging viruses, we can use de novo sequencing technology. This technology can accurately analyze the unique protein peptide sequence left by the virus in urine samples, so as to provide us with the molecular fingerprint of the new virus. Based on these sequence data, we can build a database specifically for viruses in urine samples. This database will greatly facilitate researchers and clinicians, enabling them to quickly screen out potential virus information from urine samples and achieve rapid and accurate diagnosis. In addition, the establishment of the database will also provide valuable resources for virological research and clinical treatment, and help us better understand the transmission mechanism, pathogenicity and possible intervention strategies of the virus.

